# Fourth dose of Microneedle Array Patch of SARS-CoV-2 S1 Protein Subunit Vaccine Elicits Robust Long-lasting Humoral Responses in mice

**DOI:** 10.1101/2023.10.05.561047

**Authors:** Eun Kim, Juyeop Shin, Alessandro Ferrari, Shaohua Huang, Eunjin An, Donghoon Han, Muhammad S. Khan, Thomas W. Kenniston, Irene Cassaniti, Fausto Baldanti, Dohyeon Jeong, Andrea Gambotto

## Abstract

The COVID-19 pandemic has underscored the pressing need for safe and effective booster vaccines, particularly in considering the emergence of new SARS-CoV-2 variants and addressing vaccine distribution inequalities. Dissolving microneedle array patches (MAP) offer a promising delivery method, enhancing immunogenicity and improving accessibility through the skin’s immune potential. In this study, we evaluated a microneedle array patch-based S1 subunit protein COVID-19 vaccine candidate, which comprised a bivalent formulation targeting the Wuhan and Beta variant alongside a monovalent Delta variant spike proteins in a murine model. Notably, the second boost of homologous bivalent MAP-S1(WU+Beta) induced a 15.7-fold increase in IgG endpoint titer, while the third boost of heterologous MAP-S1RS09Delta yielded a more modest 1.6-fold increase. Importantly, this study demonstrated that the administration of four doses of the MAP vaccine induced robust and long-lasting immune responses, persisting for at least 80 weeks. These immune responses encompassed various IgG isotypes and remained statistically significant for one year. Furthermore, neutralizing antibodies against multiple SARS-CoV-2 variants were generated, with comparable responses observed against the Omicron variant. Overall, these findings emphasize the potential of MAP-based vaccines as a promising strategy to combat the evolving landscape of COVID-19 and to deliver a safe and effective booster vaccine worldwide.

## Introduction

The global impact of the severe acute respiratory syndrome coronavirus 2 (SARS-CoV-2) infection has been staggering. Over 770 million confirmed cases of COVID-19 have been reported worldwide, with nearly 7 million lives lost, including more than 1 million deaths in the United States (1, 2) (until September 13, 2023). While vaccination efforts have played a crucial role in combating the pandemic, several challenges persist. Booster doses of existing vaccines have proven highly effective in preventing severe disease and fatalities. However, the virus continues to mutate, giving rise to new variants of concern (VOCs), such as the infamous Omicron and XBB subvariants. These variants have the potential to evade the immunity conferred by earlier vaccinations, raising concern about the development of updated booster shots to address the evolving threat.

One critical issue during the pandemic is the unequal distribution of vaccines globally. Many low to middle-income countries have struggled to secure adequate vaccine supplies (3, 4), leaving them without access to variant-specific vaccines that are better suited to the emerging SARS-CoV-2 landscape. Furthermore, up to 30% of the vial COVID-19 vaccines were wasted in many countries after more than a year of distribution (5). In response to these challenges, alternative devices like dissolved microneedles array patch (MAP) for subunit vaccine delivery are being developed and considered. MAP delivery of vaccines offers advantages such as dose sparing, reduced needle-stick injuries and disease transmission, resulting in a painless self-administered injection and reduced needle phobia in patients (3, 6–8). Additionally, MAPs can be pre-formulated and stably stored for extended periods at room temperature (RT), enabling distribution worldwide, including developing countries with limited cold chain supply networks (9, 10). MAP-based intradermal delivery has shown promising results in improving vaccine immunogenicity and safety in a number of vaccines studies of SARS-CoV-2 (11–15) and have been the subject of phase 1 clinical trials of influenza and Japanese encephalitis vaccination (7, 16, 17). Dissolving MAP of an HIV subunit vaccination enhanced B cell responses and significantly superior humoral immunity with a 16 and ∼ 1,300 fold elevation in bone marrow plasma cells and serum IgG titers, respectively in comparison to the traditional bolus delivery mode (18). The skin is an immunologically active tissue and contains a variety of immune cells, including antigen-presenting dendritic cells and Langerhans cells, which play a pivotal role in the body’s immune response (19). These cells can efficiently transport antigens via lymphatic drainage to initiate antigen-specific adaptive immune responses (20–22).

The spike (S) protein of SARS-CoV-2, comprising S1 and S2 subunits, mediates viral entry into host cells during infection, making it a prime vaccine target against COVID-19 (23, 24). Studies have demonstrated the MAP-based delivery of the Receptor Binding Domain (RBD) subunit vaccine induced significant B-cell and T-cell responses against S-RBD, comparable to conventional bolus injections in mice (12). Similarly, MAP have been employed to deliver both the SARS-CoV-2 S1 subunit antigen and the Toll-Like Receptor 3 (TLR3) agonist Poly(I:C). This approach has yielded robust antibody and cellular immune responses, both systemically and in the respiratory mucosa, demonstrating improved immunogenicity and safety compared to traditional intramuscular vaccines (11). Furthermore, our previous research has shown that a SARS-CoV-2 S1 subunit vaccine delivered via a MAP can rapidly stimulate the immune system to produce antigen-specific antibody responses in mice (25). In the context of ongoing research, the immunogenicity of second and third boosts with monovalent or bivalent variant vaccines from MAP-based platforms targeting wild-type, Beta, and Delta S1 antigens has been investigated. The findings indicate that MAP delivery is not only a safe but also a promising strategy for COVID-19 immunization. Second and third boosts elicited high antibody responses against S1 and efficiently neutralized the virus. Significantly, additional booster doses of MAP vaccine induced a long-lasting antibody response at high levels, persisting for at least 80 weeks. This longevity in antibody response is a promising development, as it suggests that MAP-based vaccines could provide durable protection against SARS-CoV-2 and its variants. The optimization of variant-specific MAP boosters becomes essential to booster vaccine effectiveness against current and future pandemics.

## Results

### Characterization of MAP-rS1(WU+Beta)

In our previous study, we had developed a MAP of SARS-CoV-2 S1 Wuhan (WU) subunit vaccine using CMC-based platform (25). In this study, we generated the recombinant proteins of SARS-CoV-2 S1WU, S1Beta, and S1RS09Delta and fabricated MAP using a hyaluronic acid-based droplet extension technique (DEN), which offers the benefits of being faster, scalable, and cost-effective method. The DEN MAP features a circular design with a diameter of 1.5 cm, housing a grid of 5×5 square microneedle arrays, each measuring approximately 700 μm in length.

To produce recombinant proteins of SARS-CoV-2-S1, pAd/S1Wu, pAd/S1Beta, and pAd/S1RS09Delta were generated by subcloning the codon-optimized SARS-CoV-2-S1 gene with a C-tag (EPEA) into the shuttle vector pAd (GenBank U62024) at *Sal* I and *Not* I sites (**Fig. 1A**). Variant-specific mutations for SARS-CoV-2 Beta (B.1.351) and Delta (B.1.617.2) S1 proteins are outlined. For the expression of SARS-CoV-2-S1 subunit proteins, Expi293 cells were transfected with the plasmids pAd/S1Wu, pAd/S1Beta, and pAd/S1RS09Delta. Five days after transfection, the supernatants of Expi293 cells were collected, and the recombinant proteins, rS1WU, rS1Beta, and rS1RS09Delta, were purified using C-tagXL affinity matrix and confirmed by silver staining (**Fig. 1B**). Each S1 recombinant protein was recognized by a polyclonal anti-spike of SARS-CoV-2 antibody at the expected glycosylated monomeric molecular weights of about 110 kDa under the denaturing reduced conditions. The MAP consisted of a double layered form with a base layer measuring 900 μm in base width and 200 ± 20 μm in height, and an upper layer measuring 500 ± 20 μm in length. Each MAP occupied an area size of 1.76 cm^2^, with 5 × 5 arrays of approximately 700 ± 50 μm total length. The mechanical strength (fracture force) of the MAP was measured to be more than 0.87 N, which is the required strength to penetrate the skin. 2.5 μg rS1WU plus 2.5 μg rS1Beta, or 5 μg rS1RS09Delta, were loaded in the 5×5 MAP, referred to MAP-rS1(WU+Beta) and MAP-rS1RS09Delta, respectively (**Fig. 1C and D**). To assess the delivery efficiency of the MAP, patches were reconstituted in a buffer through shaking for 30 minutes after inoculation. The concentration of remaining S1 in the patches was determined using a sandwich enzyme-linked immunosorbent assay (ELISA) employing monoclonal antibodies against SARS-CoV-2-S1 Wuhan. The findings revealed an average MAP delivery efficiency of 70% (**Fig. 1E**).

**Fig. 1.**
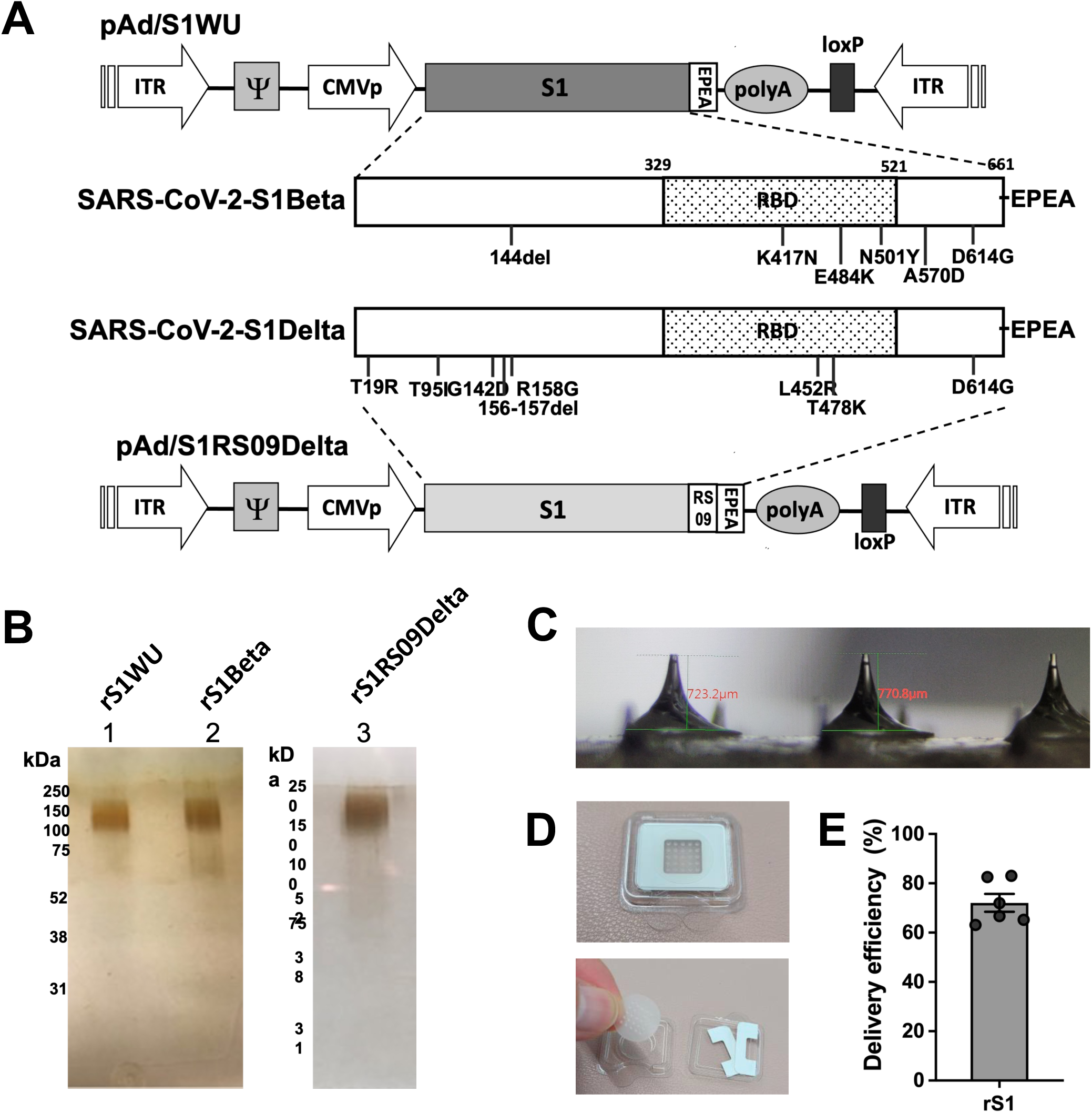
Fabrication of dissolving microneedle array patch (MAP) (A) A shuttle vector carrying the codon-optimized two variants of SARS-CoV-2-S1 gene encoding N-terminal 1-661 with a c-tag (EPEA) was designated as shown in the diagram. Amino acid changes in the SARS-CoV-2-S1 region of in this study are shown. ITR: inverted terminal repeat; RBD: receptor binding domain. (B) Silver stained SDS-PAGE of purified SARS-CoV-2 S1 proteins from Wuhan strain (lane1), Beta strain (lane 2), and S1RS09 of Delta strain (lane 3) (C and D) Image of MAP. Each MAP was approximately 700 µm in length, occupying a 5 × 5 array, 1.76 cm^2^ circle area size of the patch with rS1WU and rS1Beta (E) Delivery efficiency of rS1 by MAP into mouse skin. Recovered protein was analyzed via sandwich ELISA with S1-specific monoclonal antibody pair. Percent recovery is relative to the original control prepared fresh on the day of each assay, calculated as [(initial dose−residual dose)/initial dose x 100%].

### Enhanced Immune Response after the 2^nd^ boost with MAP-rS1(WU+Beta)

In an earlier investigation, we showed the effectiveness of a prime-boost SARS-CoV-2 S1 subunit MAP vaccine in triggering specific antibody responses in mice (25). To evaluate the influence of an additional MAP boost, we immunized BALB/c mice (5 per group) with MAP-rS1(WU+Beta) in a prime-boost regimen, spaced three weeks apart. Furthermore, we administered a supplementary boost of the identical homologous antigen intradermally at week 12 post-initial vaccination (**Fig. 2A**). Blood samples were collected before the initial immunization and every three weeks afterward, and then examined for the presence of antibodies specific to SARS-CoV-2-S1 using ELISA. As illustrated in **Fig. 2B**, the levels of serum IgG induced after the first immunization significantly surged following the booster dose, surpassing the pre-immunization levels. This indicated a robust immunogenic response, mirroring the promising outcomes observed in the prior experiment. Interestingly, the S1-specific IgG endpoint titer (EPT) experienced a rapid increase after the second booster shot and remained elevated for at least 12 weeks, gradually declining from week 15 post-second booster. However, even at week 31 post-initial vaccination, the IgG endpoint titer (EPT) remained substantially elevated in comparison to pre-immunization levels (p < 0.01, Mann-Whitney test). These findings indicate that a second booster immunization with MAP-rS1(WU+Beta) can produce strong and enduring S1-specific antibody responses, surpassing those seen after the initial booster dose.

**Fig. 2.**
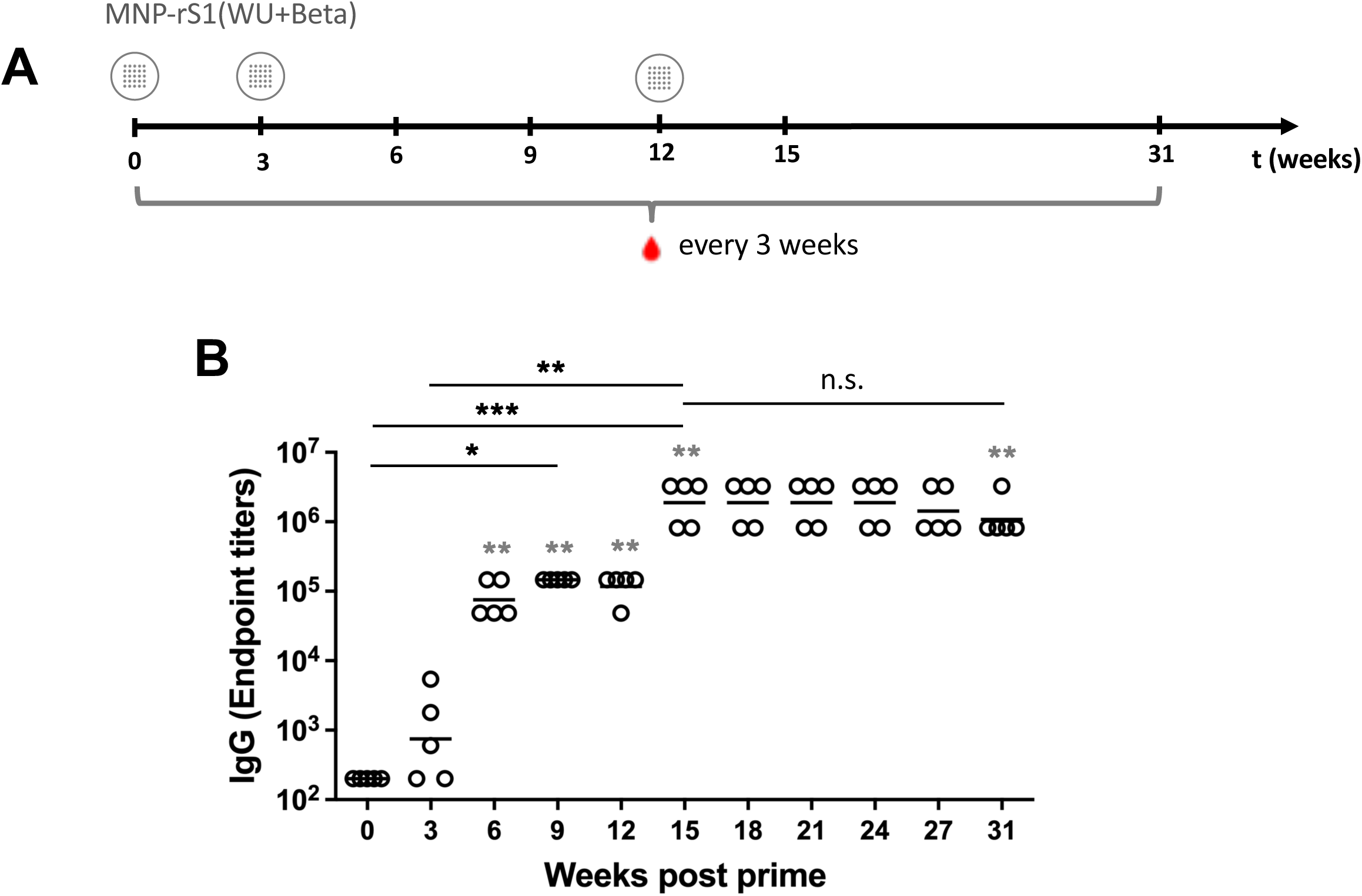
Antigen-specific antibody responses in mouse immunized with MAP of SARS-CoV-2 rS1 protein subunit vaccine after the 2^nd^ booster. (A) Schedule of immunization and blood sampling for IgG end point titration. BALB/c (N=5) were immunized with MAP-rS1(WU+Beta) at week 0, 3, and 12. The red drops denote times at which blood was drawn. (B) Sera were diluted, and SARS-CoV-2-S1-specific antibodies were quantified by ELISA to determine the IgG endpoint titer. The IgG titers at each time points were showed for each mouse. The bars represent the geometric mean. Statistical comparisons with the pre-immunized sera for ELISA were determined by an unpaired nonparametric t-test (Mann-Whitney test, grey asterisks). Differences among more than two groups were compared by the Kruskal-Wallis test, followed by Dunn’s multiple comparisons (black asterisks). Significant differences are indicated by *p < 0.05, **p < 0.01, ***p < 0.001, n.s., not significant.

### A Third Heterologous Boost of MAP Vaccine Induces Potent and Long-lasting SARS-CoV-S1 Specific Antibody Responses

To further assess the effect of an additional heterologous third boost of MAP-delivered vaccine, we generated the recombinant rS1Delta subunit vaccine which include TRL4 agonist RS09 (APPHALS) motif (25, 26) as shown in **Fig. 1**. Mice were first immunized with MAP-rS1(WU+Beta) during weeks 0, 3, and 12, and then received an additional boost of 5 μg of MAP-rS1RS09Delta at week 31 after the initial immunization. Serum samples were collected every 3 or 4 weeks until week 111 post-initial vaccination, marking the maximum duration of the study so far (**Fig. 3A**). Interestingly, following the third boost at week 31, the S1WU-specific IgG endpoint titer (EPT) didn’t increase as significantly as after the second boost. However, it remained significantly high even at 83 weeks, which marks one year after the third boost, when compared to pre-immunized sera (p < 0.01, Mann-Whitney test) (**Fig. 3B**). The fold changes of IgG EPT were, on average, 8.2-fold post-initial vaccination, 361.8-fold post-first boost, 15.7-fold post-second boost, and 1.6-fold post-third boost, respectively (**Fig. 3D**). We conducted S1Delta-specific ELISA to investigate the modest increase in IgG levels after the heterologous third boost. As depicted in Fig. 3C, following the third boost, S1Delta-specific IgG endpoint titer (EPT) did not rise as significantly as after the second boost but remained consistently high for a year, comparable to levels observed at week 34 (p < 0.01, Mann-Whitney test) (Fig. 3C). While the S1WU-specific IgG EPT did not surge as much as after the second boost, IgG levels remained consistently high for 40 weeks, starting to decline at week 46 post-third boost. Nevertheless, they still maintained significantly elevated IgG EPT even at 80 weeks post-third boost when compared to pre-immunized sera (p < 0.01, Mann-Whitney test) (**Fig. 3E**). These findings suggest that intradermal delivery via MAP induces potent and enduring SARS-CoV-S1-specific antibody responses, prolonging immune responses through an immunogenic boost.

**Fig. 3.**
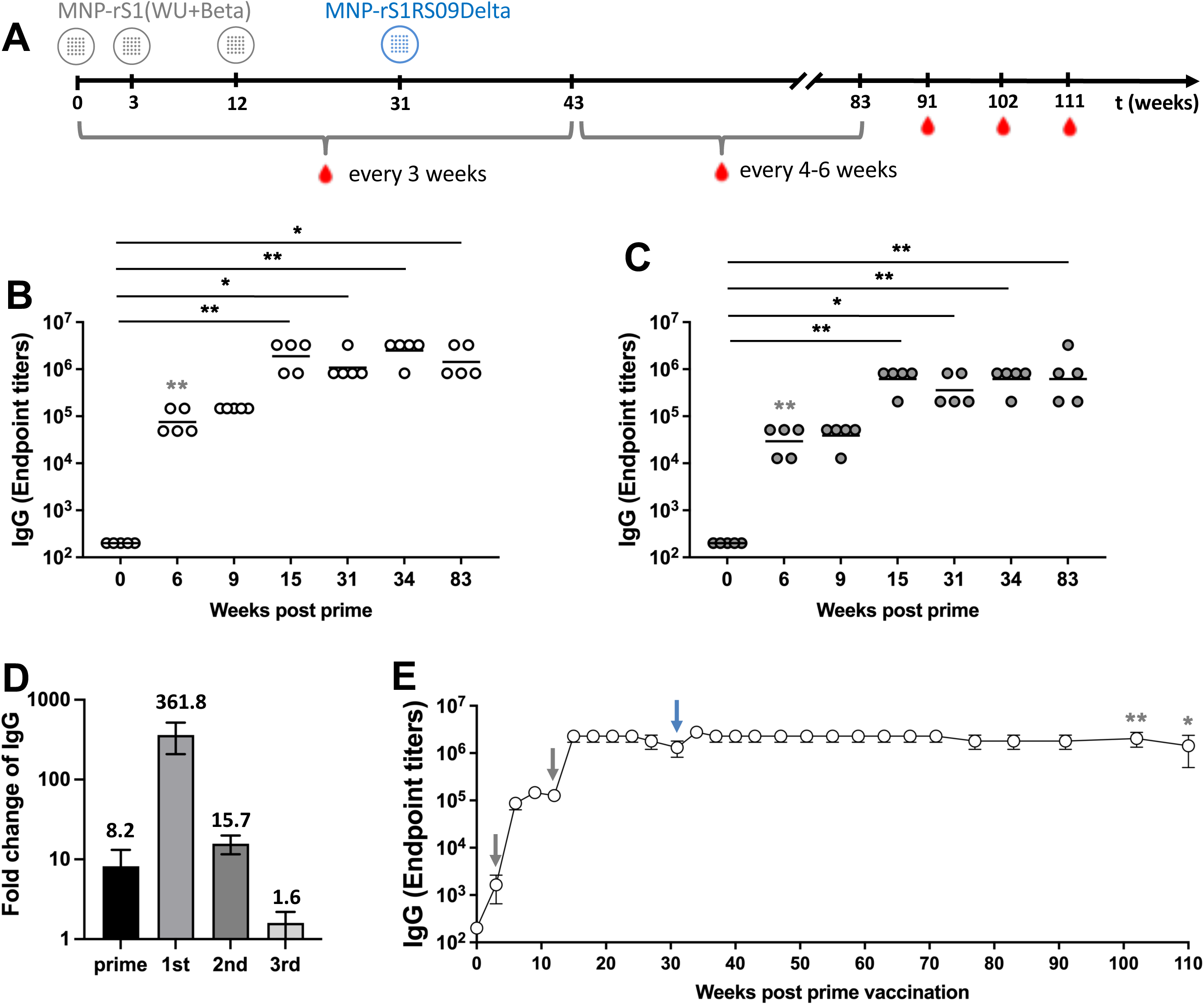
Long-lasting immune response after the 3^rd^ boost. (A) Schedule of immunization and blood sampling for IgG end point titration. BALB/c (N=5) were immunized with MAP-rS1(WU+Beta) at week 0, 3, and 12 and MAP-rS1RS09Delta at week 31. The red drops denote times at which blood was drawn. Sera at weeks 0, 6, 9, 15, 31, 34, and 83 were diluted, and SARS-CoV-2-S1WU-specific (B) and SARS-CoV-2-S1Delta-specific (C) antibodies were quantified by ELISA to determine the IgG endpoint titer. The IgG titers at each time points were showed in each mouse. The bars represent geometric mean. (D) Fold change of IgG endpoint titer after prime (W3/W0), 1^st^ (W9/W3), 2^nd^ (W15/W9), and 3^rd^ (W34/W15) boost. (E) The SARS-CoV-2-S1WU-specific IgG titers at all time points until week 111 post-prime (BALB/c (N=5) except at week 102 (N=4) and at week 111 (N=3)). Arrows along the X-axis illustrate the time points of boost vaccinations (grey arrows; MAP-rS1(WU+Beta), blue arrow; MAP-rS1RS09Delta). Statistical comparisons with pre-immunized sera for ELISA were determined by unpaired nonparametric t-test (Mann-Whitney test, grey asterisks). Differences among more than two groups were compared by Kruskal-Wallis test, followed by Dunn’s multiple comparisons (black asterisks). Significant differences are indicated by *p < 0.05, **p < 0.01.

### Distinct IgG Subtypes Levels Post-Booster Vaccination

We also determined the of S1WU-specific IgG isotypes switch, IgG1, IgG2a, IgG2b, and IgG3, following repeated booster vaccinations, indicating a prevalent and/or balanced Th2-or Th1-like immune response (**Fig. 4**). Sera collected at weeks 0, 9, 15, 34, and 83 post-prime were subjected to isotype-specific ELISA. All subclasses showed a significant increase after the 2^nd^ boost and maintained significantly high levels until weeks 83, in comparison to pre-immunized sera, except for IgG3, which exhibited a lower level among the IgG subclasses. Interestingly, the immune responses switched toward a Th1 bias response at week 15 (post-2^nd^ boost) and displayed a less varied Th2/Th1 ratio among the immunized mice compared to those at weeks 9 (post-1^st^ boost) (SD=6.89 ± 3.08 at week 9 vs. SD=2.09 ± 0.94 at week 15). The geometric mean titer (GMT) of IgG1 and IgG2b increased slightly after the 3^rd^ boost, while those of IgG2a and IgG3 was not increased. Thus, these results suggest that a Th2-prevalent responses were observed at all time points based on the ratio of IgG1 to IgG2a + IgG2b + IgG3 antibody subclasses. Furthermore, it was evident that at least three doses of MAP were required to induce significantly high IgG subclasses antibody titers compared to pre-immunized sera.

**Fig. 4.**
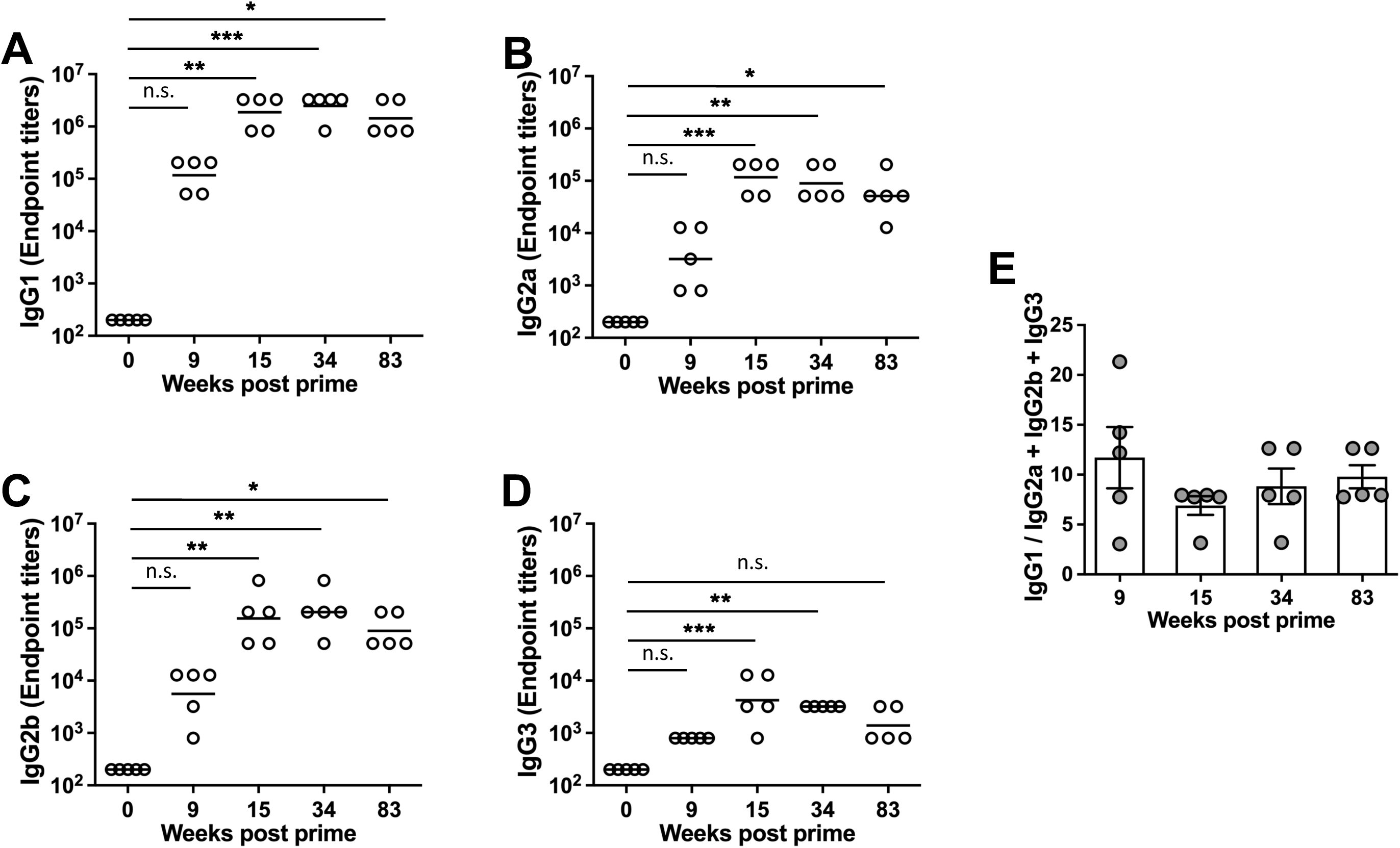
IgG subclasses after boosters. BALB/c (N=5) were immunized with MAP-rS1(WU+Beta) at week 0, 3, and 12, and MAP-rS1RS09Delta at week 31. Sera at weeks 0, 6, 9, 15, 31, 34, and 83 were diluted, and SARS-CoV-2-S1WU-specific IgG1 (A), IgG2a (B), IgG2b (C), and IgG3 (D) were quantified by ELISA to determine each IgG subclasses endpoint titer. The titers at each time points were showed for each mouse. The bars represent geometric mean. (E) S1-specific IgG1/IgG2a+IgG2b+IgG3 ratios of individual mice at weeks 9, 15, 34, and 83 as mean values with SEM. Groups were compared by the Kruskal-Wallis test at each time point, followed by Dunn’s multiple comparisons. Significant differences are indicated by *p < 0.05, **p < 0.01, ***p < 0.001, n.s., not significant.

### Neutralizing Antibody Levels after the Boosters

To evaluate the presence of SARS-CoV-2-specific neutralizing antibodies generated by repeated boosters, we performed a microneutralization assay (VNT_90_) to assess the ability of sera from immunized mice to neutralize the infectivity of SARS-CoV-2 variants, including Wuhan, Delta (B.1.617.2), and Omicron (BA.1) variants (**Fig. 5**). SARS-CoV-2-neutralizing antibodies were under the limit of detection at week 9, after the first boost. However, sera from one to three out of 5 mice were shown the neutralization activity at week 15 following the second boost. Interestingly, neutralizing antibodies against not only Wuhan and Delta but also the Omicron variant were generated, with comparable responses after three and fourth doses. The GMT of VNT_90_ at weeks 15, 34, and 83 were as follows: 7.6, 11.5 and 15.2 against Wuhan; 8.7, 15.2 and 17.4 against Delta; 10.0, 10.0 and 11.5 against Omicron, respectively. Interestingly, only Wuhan and Delta VNT_90_ at weeks 83 were statistically significant compared to those at week 0, with no significant differences observed against Omicron. The fold change of GMT of VNT_90_ against Wuhan, Delta, and Omicron at weeks 83 (at 52 weeks post-3^rd^ boost) compared to those at week 34 (at 2 weeks post-3^rd^ boost) was 1.32-fold, 1.15-fold, and 1.15-fold, respectively. These findings unmistakably demonstrate that administering a minimum of three doses of MAP can generate detectable neutralizing antibodies against various SARS-CoV-2 variants, including Omicron (BA.1).

**Fig. 5.**
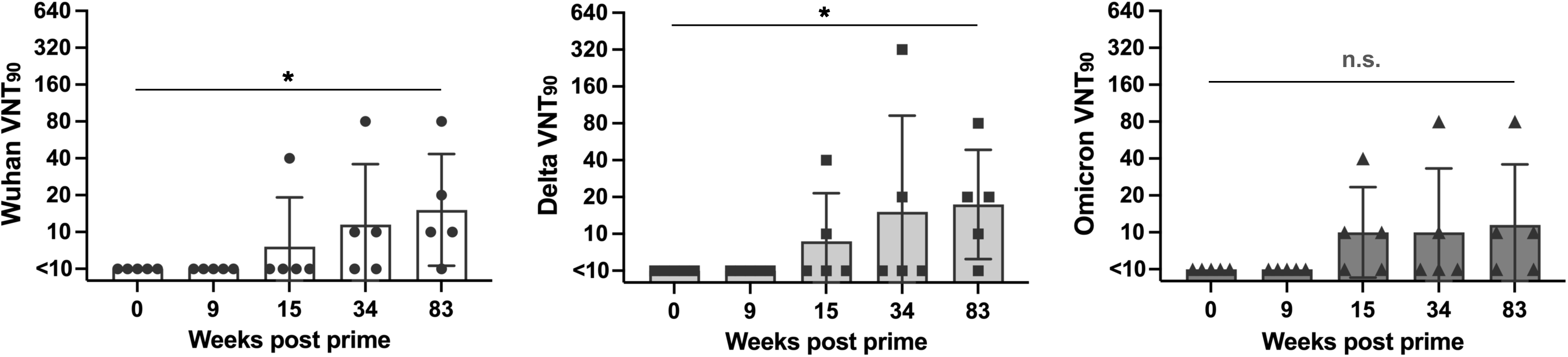
Neutralization of SARS-CoV-2 variants. Serum from mice immunized with SARSCoV-2 S1 via MAP intradermal delivery was assessed using a microneutralization assay (VNT_90_) for neutralization against SARS-CoV-2 variants (A) Wuhan, (B) Delta variant (B.1.617.2), and (C) Omicron variant (BA.1). Data are representative of the geometric mean with error bars representing geometric standard deviation. Groups were compared by the Kruskal-Wallis test at each time point, followed by Dunn’s multiple comparisons. Significant differences are indicated by *p < 0.05, n.s., not significant.

## DISCUSSION

In our prior research, we established that a SARS-CoV-2 S1 subunit MAP vaccine swiftly and effectively triggered the immune system, generating specific antibody responses in mice (25). In this current study, we examined the immunogenicity of the second and third boosts utilizing monovalent and bivalent variant vaccines delivered through MAP-based platforms, focusing on wild-type, Beta, and Delta S1 antigens. Our findings revealed that the second and third boosts elicited robust antibody responses against S1, effectively neutralizing the virus. Importantly, these additional MAP vaccine boosts resulted in enduring high-level antibody responses, persisting for at least 80 weeks.

In our previous work, we utilized MAP made from CMC (carboxymethyl cellulose) using a two-step spin-drying technique through micro-molding. CMC is a biocompatible, biodegradable polymer commonly employed to enhance solution viscosity for MAP materials (27, 28). However, in this study, we opted for hyaluronic acid (HA), a naturally occurring polymer found in humans and animals with no adverse effects (29), as the MAP material (Fig. 1). HA is continuously secreted by various specialized cells in the body and is naturally broken down by enzymes (hyaluronidases) (30). The dissolution rates of MAP in the skin vary based on MAP dimensions, shape, and animal species (31, 32). As a result, both CMC and HA-based MAPs completely dissolved in rat skin within 5 minutes and in human skin within 2 hours, respectively. Notably, fluorescein-labeled ovalbumin released from HA-based MAP diffused more rapidly than from CMC-based MAP and persisted in rat skin for a longer duration (48 hours vs. 24 hours) (33), demonstrating an adjuvant function. Indeed, HA acts as an immunological adjuvant for protein-based vaccines (34, 35). The chemical bonding of HA to antigens stimulated strong and enduring humoral responses without requiring additional immunostimulatory compounds, leading to reduced booster needs and enabling antigen dose conservation (35). Furthermore, in this study, we employed the DEN fabrication method, which outperforms the previous micro-molding technique in terms of speed and ease of mass production by eliminating multiple steps, including the creation of a master mold (36). This significant advancement underscores the potential of DEN MAP technologies as an advanced and promising approach for future clinical production in global mass vaccination efforts.

In this study, we found that administering a second boost of bivalent MAP-S1(Wu+Beta) resulted in a substantial 15.7-fold increase in IgG GMT (**Fig. 2**). This boost led to neutralizing activity in the sera of some mice (**Fig. 5**), suggesting its efficacy in conferring protection against viral infection. Following a third boost with the monovalent MAP-S1RS09Delta, there was only a modest 1.6-fold increase in IgG GMT between the third and fourth doses (**Fig. 3**). This suggests that the vaccine had already reached a high level, and further increases were limited. This phenomenon might be attributed to the loss of short-lived plasma cells in the early stages, causing the initial rapid decline in S1-specific antibodies during the prime-boost. Conversely, the development of long-lived memory plasma cells with additional boosts could explain the plateau in the antibody response (37). The relatively modest increase in IgG levels between the third and fourth doses could be attributed to the short interval, which might not provide sufficient time for the IgG titer to decline. Research on healthcare workers who received the Pfizer mRNA COVID19 vaccine revealed that neutralizing antibody concentrations were higher when the second vaccine dose was administered within an interval of 6-14 weeks, compared to the recommended 3–4-week regimen (38). Similarly, the approach of delaying the second doses of mRNA vaccination may enhance humoral immune responses, including improved virus neutralization against both wild-type and variant SARS-CoV-2 viruses (39). Allowing ample time for processes such as antibody somatic mutation, turnover of memory B cell clones, ongoing B-cell clonal selection, and the accumulation of monoclonal neutralizing antibodies that display exceptional resistance to SARS-CoV-2 RBD mutations, including those found in VOCs (Variants of Concern), could be crucial (40). Notably, only the Wuhan and Delta VNT90 at week 83 showed statistical significance compared to those at week 0, with no significant differences observed at week 34, which was 2 weeks after the third boost (Fig. 5). Moreover, following the second and third boosts, neutralizing antibodies against the Omicron variant were induced at comparable levels to those against Wuhan and Delta, the strains chosen for subunit vaccination in this study.

Nevertheless, our research revealed that even after the fourth vaccine dose, IgG levels remained consistently elevated for over 111 weeks (**Fig. 3E**). Past studies have shown that vaccine-induced protection against SARS-CoV-2 diminishes over time, leading to instances of SARS-CoV-2 reinfection (41–43). Consequently, the challenge of sustaining long-term immunity is the major obstacles in enhancing and approving COVID-19 vaccines (44). While it’s complex to directly compare human and mouse lifespans, the enduring immune responses observed in this study following additional MAP boosts suggest a potential solution to overcome these hurdles. The third dose of MAP-S1 triggered statistically significant IgG1, IgG2a, IgG2b, and IgG3 subclasses specific to S1, revealing a balanced Th1/Th2 immune response with a slight Th2 prevalence (**Fig. 4**) during the timeframes we investigated. Following the third boost, IgG1 and IgG2b GMTs showed a slight increase, whereas IgG2a and IgG3 levels remained unchanged. In humans, IgG4 production is closely linked to persisting antigens (45). Indeed, noninflammatory IgG4 levels rose several months after the second and third BNT162b2 mRNA vaccinations (46). This class switch was associated with reduced spike-specific antibody capabilities to mediate crucial antiviral immune functions, such as antibody-dependent cellular phagocytosis (ADCP) and antibody-dependent complement deposition (ADCD) (47). Although murine IgG subclasses differ from human counterparts, human IgG4 functions similarly to murine IgG1, displaying anti-inflammatory properties without complement fixation, and engaging with inhibitory FcγRIIb (46). It’s crucial to highlight that the IgG2a levels remained statistically significant until week 83, playing a vital role in antiviral immunity through ADCP and ADCD, owing to its strong FcγR-mediated activity and complement-fixing functions. To steer the IgG subclass toward a Th1-biased immune response, incorporating vaccine adjuvants is anticipated. Several studies have shown that adjuvanted SARS-CoV-2 MAP vaccines induce a balanced Th1/Th2 response in mice (11, 48, 49). Additionally, incorporating vaccine adjuvants can boost vaccine efficacy and conserve more vaccine doses (50). Nevertheless, adjuvanted intradermal vaccines carry a risk of causing significant local reactions. Intradermal vaccine delivery without adjuvants results in more frequent and severe local adverse reactions compared to intramuscular vaccination (51, 52). This can be explained by the presence of immune cells in the skin, such as LCs, DCs, and macrophages, which promote both adaptive immunity and local adverse reactions by releasing cytokines and chemokines, as well as recruiting peripheral neutrophils, monocytes, and eosinophils. Therefore, adjuvants for intradermal MAP vaccination might ensure better local safety compared to those used in intramuscular vaccination (11, 53). Similarly, an adjuvanted MAP with a subunit booster strategy is likely to have a beneficial effect on protection, particularly against distant variants such as Omicron XBB subvariants. Indeed, MAPs containing both SARS-CoV-2 S1 subunit antigen and the TLR3 agonist Poly(I:C) induced long-lasting antibodies compared to S1 MAP without an adjuvant (11). High-density microarray patch (HD-MAP)–delivered spike with the saponin-based adjuvant QS21 resulted in a robust neutralizing IgG and cellular immune response in mice, surpassing the same vaccine without an adjuvant. This approach provided complete protection from a lethal SARS-CoV-2 challenge after only one dose, with no virus replication observed in the lungs or brain when applied to K18-hACE2 mice (13). Moreover, adjuvants serve as a suitable strategy to direct the adaptive immune response toward the desired T helper subset, such as Th1, which is dominant for protection against viral diseases (11, 54).

This study had two limitations, which will be addressed in future research. These include the assessment of T-cell specific cellular immune responses and conducting a SARS-CoV-2 challenge study to evaluate the protective effectiveness of a booster vaccination. Nevertheless, previous studies have shown that intradermal delivery of SARS-CoV-2 spike protein using dissolvable microneedle patches activated T-cell immunity (11, 12, 48). MAP S-RBD elicited significant B-cell and T-cell responses against S-RBD, comparable to conventional bolus injection (12).

In summary, our research investigated the impact of second and third boosts using MAP-based platforms targeting SARS-CoV-2 S1 antigens, resulting in the development of strong and enduring antibody responses. These findings have implications for further research on using MAP to deliver S1 subunit vaccines for emerging SARS-CoV-2 variants, potentially serving as boosters to enhance cross-neutralizing antibodies. This approach could offer a viable platform for widespread global vaccination efforts.

## Materials and methods

### Recombinant Proteins Expression and Purification

Codon-optimized cDNA encoding the Wuhan or Beta SARS-CoV-2-S1 glycoprotein, encompassing amino acids 1 to 661 (25, 55, 56) with C-tag was synthesized and cloned into pAdlox between the *Sal* I and *Not* I sites. Similarly, for codon-optimized Delta (B.1.617.2) SARS-CoV-2-S1 glycoprotein, spanning amino acids 1 to 661 and equipped with the RS09 TLR4 agonist and C-tag, synthesis and cloning into pAdlox were performed using the same method. The production of rS1WU, rS1Beta, and rS1RS09Delta involved transient expression in Expi293 cells with pAd/S1WU, pAd/S1Beta, and pAd/S1RS09Delta, respectively, utilizing the ExpiFectamie^TM^ 293 Transfection Kit (ThermoFisher) as previously reported (56, 57). Subsequently, the recombinant proteins were purified using a CaptureSelect^TM^ C-tagXL Affinity Matrix prepacked column (ThermoFisher), followed the manufacturer’s guidelines as previously detailed (56, 57). In brief, the C-tagXL column underwent conditioning with 10 column volumes (CV) of equilibrate/wash buffer (20 mM Tris, pH 7.4) before sample application. The supernatant was adjusted to 20 mM Tris with 200 mM Tris (pH 7.4) before loading onto a 5-mL prepacked column at a rate of 5 ml/min. The column underwent subsequent washing cycles, alternating between 10 CV of equilibrate/wash buffer, 10 CV of strong wash buffer (20 mM Tris, 1 M NaCl, 0.05% Tween-20, pH 7.4), and 5 CV of equilibrate/wash buffer. The recombinant proteins were eluted from the column using an elution buffer (20 mM Tris, 2 M MgCl_2_, pH 7.4). The eluted solution was desalted and concentrated with preservative buffer (PBS) in an Amicon Ultra centrifugal filter device with a 50,000 molecular weight cutoff (Millipore). The concentration of the purified recombinant proteins was determined by the BCA protein assay kit (Thermo Scientific) with bovine serum albumin (BSA) as a protein standard. The proteins were separated by reducing SDS-PAGE and visualized through silver staining.

### Preparation of Dissolving Microneedle Array Patches

Dissolving microneedle array patch (MAP) containing the protein rS1WU combined with rS1Beta or rS1RS09Delta were fabricated using the DEN (droplet extension technique) method. In this technique, droplets were solidified and formed into cone shape microstructure by air blowing, as previously descried (58). The MAP used in this study was provided by Raphas Co., Ltd. (Rep. of Korea). Briefly, a 5 x 5 base array was dispensed onto a pattern-mask hydrocolloid adhesive sheet (Hiks C&T) using a customized MPP-1 dispenser (Musashi) with 10% hyaluronic acid (HA, Kikkoman). The dispensed array was then dried overnight at room temperature. Subsequently, a viscous solution containing the antigens and HA (Contipro Inc) was dispensed onto a different base array. The two dispensed droplets were brought into contact and extended to the target length, and then symmetric air blow was applied at room temperature to solidify the extended viscous droplets, forming cone-shaped microstructures. The mechanical strength of each MAP was measured using the universal test machine (UTM, ZwickRoell). Briefly, a MAP was mounted on the UTM stage and aligned with the UTM probe. The probe was moved vertically until the microneedle fractured, with the force responsible for breaking the microneedle recorded as its mechanical strength.

### Quantification of Release from MAP

To reconstitute the MAP, 1 ml of PBS was added to each well of a 12-well plate containing the MAP. Reconstitution was achieved by pipetting and incubating for 30 min at RT with shaking at 700 rpm. Following a 10-fold dilution of the collected solution, the delivered rS1 in the patches was quantified using a sandwich ELISA, employing SARS-CoV-2 S1-specific monoclonal antibody pairs, as previously described (56). The amount of protein in a sample was determined using a curve generated from purified SARS-CoV-2 S1 standards for known concentration, The results were calculated as follows: Delivery efficiency of rS1 on MAP (%) =L100 × (1 − (rS1 concentration of the collected patch after inoculation / rS1 concentration of the origin patch))

### Animal immunization

MAPs loaded with of 2.5 µg of rS1WU and 2.5 µg of rS1Beta, referred to as MAP-rS1(WU+Bata), were applied to the skin of the back region of female BALB/c mice (n = 5 animals per group). Prior to vaccination, the hair at the vaccination site was removed by shaving and depilatory cream. MAP was applied to the dehaired back skin of a mouse using a handheld spring applicator and held in place by finger pressure for 10 seconds, followed by leaving it on the skin for more than 2 hours. No skin adverse effects, such as skin irritation at the vaccinated region were observed. At 3 and 12 weeks after the primary immunization, mice were received booster immunizations with homologous immunogens. Blood samples were collected from the retro-orbital vein of mice at every three weeks until weeks 31. The obtained serum samples were diluted and used to evaluate rS1WU-specific antibodies via ELISA. To assess of the effect of 3rd boost, MAPs loaded with 5ug of rS1RS09Delta (MAP-rS1RS09Delta) were inoculated intradermally at week 31 post-prime. Blood samples were collected from the retro-orbital vein at every three to ten weeks until weeks 111. The obtained serum samples were diluted and used to evaluate both rS1WU– and rS1Delta-specific antibodies by ELISA. Mice were maintained under specific pathogen-free conditions at the University of Pittsburgh, and all experiments were conducted in accordance with animal use guidelines and protocols approved by the University of Pittsburgh’s Institutional Animal Care and Use (IACUC) Committee.

### Assessment of Serum Humoral Antibodies

Serum humoral antibodies generated against spike protein were assessed using ELISA, as previously described (25, 55, 56). Sera from all mice were collected before vaccination and then every 3 or 4 weeks after immunization. These sera were tested for SARS-CoV-2 S1WU-or S1Delta-specific IgG antibodies using conventional ELISA. Furthermore, sera collected at week 0, 9, 15, 34, and 83 after vaccination were also examined for SARS-CoV-2-S1-specific IgG1, IgG2a, IgG2b, and IgG3 antibodies using ELISA. In brief, ELISA plates were coated with 200 ng of recombinant SARS-CoV-2-S1WU protein (Sino Biological) per well overnight at 4°C in carbonate coating buffer (pH 9.5) and then blocked with PBS containing 0.05% Tween 20 (PBS-T) and 2% bovine serum albumin (BSA) for one hour. Mouse sera were serially diluted in PBS-T with 1% BSA and incubated overnight. After washing the plates, anti-mouse IgG-horseradish peroxidase (HRP) (1:10000, Jackson Immunoresearch) was added to each well and incubated for one hour. For IgG1, IgG2a, IgG2b, and IgG3, HRP-conjugated IgG1, IgG2a, IgG2b, and IgG3 (1:20000, Jackson Immunoresearch) were added to each well and incubated for 1 hour. The plates were washed three times, developed with 3,3’5,5’-tetramethylbenzidine, and the reaction was stopped. Absorbance at 450 nm was determined using an ELISA reader (Molecular Devices SPECTRAmax).

### SARS-CoV-2 microneutralization assay

Neutralizing antibody titers against SARS-CoV-2 were defined according to the following protocol (59, 60). Briefly, 50 µl of sample from each mouse, starting from 1:10 in a twofold dilution, were added in two wells of a flat bottom tissue culture microtiter plate (COSTAR), mixed with an equal volume of 100 TCID_50_ of a SARS-CoV-2 Wuhan, Delta (B.1.617.2), or Omicron (BA.1) strain isolated from symptomatic patients, previously titrated. After 1 hour incubation at 33°C in 5% CO_2_, 3 x 10^4^ VERO E6 cells were added to each well. After 72 h of incubation at 33°C 5% CO_2_, wells were stained with Gram’s crystal violet solution plus 5% formaldehyde 40% m/v (Carlo ErbaSpA, Arese, Italy) for 30 min. After washing, wells were scored to evaluate the degree of cytopathic effect (CPE) compared to the virus control. Neutralizing titer was the maximum dilution with a reduction of 90% of CPE. A positive titer was equal to or greater than 1:10. The GMT of VNT_90_ endpoint titer was calculated with 5 as a negative shown <1:10. Sera from mice before vaccine administration were always included in VNT assay as a negative control.

### Statistical analysis

Statistical analyses were performed using GraphPad Prism v9 (San Diego, CA). Antibody endpoint titers and neutralization data were analyzed by Kruskal-Wallis test, followed by Dunn’s multiple comparisons. A Mann-Whitney U test was used for intergroup statistical comparison. Data were presented as the means ± standard errors of the mean (SEM) or geometric means ± geometric errors. Significant differences are indicated by * p < 0.05, **p < 0.01, ***p < 0.001.

## Notes

### Competing Interest Statement

The authors declare that they have competing interests regarding the research presented in this manuscript. Specifically, AG and EK are co-founders of GAPHAS PHARMACEUTICAL INC., a private startup company that could potentially benefit from the findings of this research. AG, EK, and MSK have equity in GAPHAS PHARMACEUTICAL INC. However, the authors have taken measures to ensure that the research is conducted objectively and that the data and conclusions presented in this manuscript are not influenced by their competing interests. The study was designed, conducted, and analyzed independently of the company. The authors also declare that GAPHAS PHARMACEUTICAL INC. did not provide financial or material support for this research.

